# Nonspecific Cleavages Arising from Reconstitution of Trypsin under Mildly Acidic Conditions

**DOI:** 10.1101/2020.05.26.116509

**Authors:** Ben Niu, Michael Martinelli, Yang Jiao, Eric Meinke, Mingyan Cao, Jihong Wang

## Abstract

Tryptic digestion of proteins followed by liquid chromatography with tandem mass spectrometry analysis is an extensively used approach in proteomics research and biopharmaceutical product characterization, owing to the high level of cleavage fidelity produced with this technique. However, nonspecific trypsin cleavages have been frequently reported and shown to be related to a number of digestion conditions and predigestion sample treatments. In this work, we reveal that, for a number of commercial trypsins, reconstitution and storage conditions can have a significant impact on the occurrence of trypsin nonspecific cleavages. We analyzed the tryptic digestion of a variety of biotherapeutics, using trypsins reconstituted under different conditions. The results indicate that, for many commercial trypsins, commonly recommended reconstitution/storage conditions (mildly acidic, e.g., 50 mM acetic acid, 1 mM HCl) can actually promote nonspecific trypsin activities, which are time dependent and can be as high as 20% in total relative abundance. In contrast, using water for reconstitution and storage can effectively limit nonspecific cleavages to 1%. Interestingly, the performances of different commercial trypsins were found to be quite distinct in their levels of nonspecific cleavages and responses to the two reconstitution conditions. Our findings demonstrate the importance of choosing the appropriate trypsin for tryptic digestion and the necessity of assessing the impact of trypsin reconstitution and storage on nonspecific cleavages. We advocate for manufacturers of commercial trypsins to reevaluate manufacturing processes and reconstitution/storage conditions to provide good cleavage specificity.

## Introduction

Owing to its ready availability and high fidelity, trypsin is by far the most widely used proteolytic enzyme in mass spectrometry (MS)–based research and applications [1–6]. These applications rely on the ability of liquid chromatography (LC) with tandem mass spectrometry (LC-MS/MS) to identify and quantify various peptide species with a high degree of accuracy, sensitivity, and reproducibility. Studies using trypsin-based digestion processes, however, have often reported nontryptic activities, which generate semitryptic and nontryptic peptides through nonspecific cleavages (cleavages at residues other than Arg or Lys) [7–10]. Although the use of nonspecific cleavage products might contribute to improved protein sequence coverage and identification [11, 12], they are also accompanied by unexpected peptides in the tryptic digestion profile. These nontryptic cleavages disperse the signals of the specifically cleaved peptides that are available for detection and increase the database searching workload for a complex sample matrix, potentially affecting accurate identification [13, 14].

Tryptic digestion–based peptide mapping is commonly used in the biopharmaceutical industry to quantitate posttranslational modifications (PTMs) of a biotherapeutic, to provide identity confirmation, and to evaluate purity [15, 16]. The unpredictable emergence of nontryptic cleavages can pose great challenges to achieving assay fidelity and reproducibility. The implementation of multi-attribute method (MAM) analysis, a more recent elaboration from tryptic peptide mapping [16–18], can also be affected, owing to the introduction of new peaks into, or removal of peaks from, the chromatographic profile. It is therefore critical to minimize nontryptic activities during trypsin-involved digestion processes.

Nontryptic activities typically indicate the presence of proteases other than trypsin, such as chymotrypsin contamination [19]. However, most commercial trypsins have been treated with *N*-*p*-tosyl-L-phenylalanyl chloromethyl ketone (TPCK) during the manufacturing process to remove traces of chymotrypsin [20]. In addition, studies have shown that trypsin cleavage specificity can be affected by a number of experimental conditions, including temperature, pH, enzyme-to-substrate ratio, and duration [7, 10, 21]. The variable quality of commercial trypsins can also have a great impact on cleavage specificity [22, 23]. In this study, however, we report that trypsin reconstitution and storage conditions alone can have a significant impact on the extent of nonspecific trypsin cleavages.

Herein we demonstrate that the reconstitution of several commercial trypsins in mildly acidic conditions (50 mM acetic acid or 1 mM HCl) can lead to dramatically increased nontryptic activities, giving rise to numerous semitryptic and nontryptic peptides. More important, although these reconstitution and storage conditions are recommended by manufacturers, the extent of nontryptic activities increases as a function of reconstitution and storage time and could be minimized for some trypsins if reconstituted and/or stored in water. To elucidate cleavage preferences for nontryptic activities, we used new peak detection (NPD) analysis to detect and identify semitryptic and nontryptic peptides and characterized the residues accountable for compromised cleavage specificity. Our work demonstrates the impact of mildly acidic reconstitution and storage conditions on nonspecific trypsin cleavages and reveals that, for several commercial trypsins, such conditions are inappropriate for trypsin reconstitution and storage. Although the integrity of the digestion profile may be compromised because of reconstitution and/or storage conditions, some trypsins manifested better specificity than others. Given these largely variable trypsin specificities, we show the necessity of assessing nonspecific cleavages, and identify several diagnostic peptides from NISTmAb digests that can be used for quick evaluation of the extent of nonspecific cleavages. We recommend that manufacturers of commercial trypsins reevaluate manufacturing processes and reconstitution/storage conditions for improved trypsin cleavage specificity.

## Materials and Methods

### Materials and chemicals

All biologics were produced and purified at AstraZeneca (Gaithersburg, MD) and were stored at –80°C at pH ~6.0 before use. Unless noted otherwise, trypsin was purchased from Promega (V5280; Madison, WI) and is denoted as Trypsin-1. Seven other trypsins were used and are denoted as Trypsin-2 (V5111; Promega), Trypsin-3 (786-254; G-Biosciences), Trypsin-4 (786-254B; G-Biosciences), Trypsin-5 (EN-151; Princeton Separations), Trypsin-6 (11418025001; Roche), Trypsin-7 (90057; Pierce), and Trypsin-8 (T-6567; Sigma-Aldrich).

Guanidium hydrochloride solution (8 M concentration), iodoacetamide (IAM), dithiothreitol, and hydrochloric acid microdialysis cassettes and devices (10 K MWCO) were purchased from Thermo Fisher Scientific (Waltham, MA). Urea powder was purchased from GE Healthcare Life Sciences (Chicago, IL). Tris HCl buffers were purchased from G-Biosciences. Acetic acid and trifluoroacetic acid (TFA) were purchased from Sigma. High-performance liquid chromatography (HPLC)–grade water and acetonitrile were purchased from Honeywell (Muskegon, MI).

### Sample preparation

Protein samples in formulation were diluted to 5.0 mg/mL with HPLC-grade water and denatured with 8.0 M guanidine HCl. Disulfide bonds were reduced with 30 mM dithiothreitol for 30 min at 37°C, followed by alkylation of free thiols with 70 mM IAM for 30 min at room temperature in the dark. Buffer exchange was conducted with microdialysis cassettes according to the manufacturer’s recommendations. Samples were buffer exchanged to a solution containing 2 M urea, 150 mM Tris at pH 7.4 for tryptic digestion.

### Tryptic digestion

All trypsins were stored in lyophilized form at temperatures recommended by the manufacturers and were brought to room temperature before use. Trypsins were reconstituted to 0.67 mg/mL with different solvents (HPLC-grade water, 1 mM HCl, 50 mM acetic acid) and held at different temperatures for various periods as needed before being used for digestion with a mass ratio of 1:12 (trypsin:protein). The digestion was incubated at 37°C for 3.5 h before being quenched by the addition of 1% TFA.

### LC-MS/MS

For LC-MS/MS, mobile phase A consisted of 0.02% TFA in HPLC-grade water and mobile phase B consisted of 0.02% TFA in HPLC-grade acetonitrile. Six micrograms of digest was loaded onto an Acquity BEH reversed-phase C18 column (130 ◻, 1.7 μm, 2.1 × 150 mm; Waters, Milford, MA). The LC was operated at a flow rate of 0.2 mL/min, with the column temperature kept at 55°C during separation and the autosampler temperature at 6°C. Total run time per sample was 90 min.

A Q Exactive HF-X mass spectrometer (Thermo Fisher Scientific), operated in positive polarity mode, was used for mass detection. The scan range of precursor ions was set at 250– 2,000 m/z for all samples, with a high mass resolving power of 120,000. Data acquisition was performed in top 5 data-dependent acquisition mode, in which the five most abundant precursor ions corresponding to peptide elution per scan were subjected to higher-energy C-trap dissociation (HCD) in the HCD cell to obtain production mass spectra. The normalized collision energy was set to 35% of maximum. Dynamic exclusion was activated for 8 s after each scan to enable MS/MS fragmentation of lower-abundance peptides. The maximum injection time was set to 150 ms for a full mass spectral scan and to 50 ms for each MS/MS scan.

### Database searching

Raw data files were subjected to database searching with Byos (Protein Metrics, San Carlos, CA) against the amino acid sequences of corresponding biologics and reversed decoys of all possible peptides [24]. The searching parameters were set to include IAM alkylation as fixed modification for Cys-containing peptides and a number of common PTMs as variable modifications, including deamidation, oxidation, N-succinimide, D-succinimide, dioxidation, pyroglutamate formation, W-kynurenine formation, and amidated proline. The search tolerance window was set to 10 ppm for all precursor ions and to 50 ppm for product spectra. The C-termini of Arg and Lys were selected as fully specific cleavage sites, with missed cleavage tolerance as 3. For database searching of nonspecific cleavages, the digestion specificity was set to “nonspecific” with any number of missed cleavages.

Relative levels of nonspecific cleavages were calculated as:

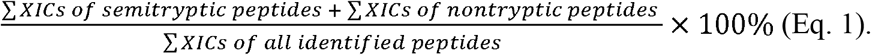

Peptides that were identified as semitryptic or nontryptic but had retention times identical to those of their corresponding fully tryptic parent peptides were considered to be from in-source fragmentation and were not included in the calculation of relative levels of nonspecific cleavages [25].

### New peak detection analysis

Progenesis QI software (Waters) with Byonic node was used for NPD analysis through binary comparison of the raw data. The absolute NPD intensity threshold was set to 1.0E6 units. The fold-change threshold, which is used to designate a peak as “new”, was set to threefold as minimum. The detected “new peaks” could be imported to Byonic (in .mgf format) for nonspecific cleavage database searching.

## Results and Discussion

### Increased nonspecific cleavages with acidic reconstitution

Most manufacturers recommend reconstituting lyophilized trypsin by using mildly acidic solutions (e.g., 50 mM acetic acid or 1 mM HCl) to sustain tryptic activities and inhibit autolysis during storage before reuse. Trypsin-1 was reconstituted in 50 mM acetic acid (termed “condition 1”) prior to digestion and the outcome was compared with that of trypsin reconstituted in HPLC-grade water (termed “condition 2”). Results showed that the digestion profiles, as represented by UV and/or total ion chromatograms (TICs), were noticeably different; multiple new peaks arose in condition 1, indicating the presence of new peptides.

To illustrate, reference material of monoclonal antibody A (mAb-A) were subjected to the tryptic digestion protocol, using identical trypsins but different reconstitution conditions. One trypsin was reconstituted in 50 mM acetic acid (condition 1), and the other was reconstituted in HPLC-grade water (condition 2). UV chromatograms of both digestions are shown in Fig 1A. The major UV peaks observed in condition 2 corresponded to the fully tryptic mAb-A heavy-chain and light-chain peptides, with a few minor peaks associated with trypsin autolysis (denoted with asterisks in Fig 1A). However, the digestion profile in condition 1 differed; in addition to the fully tryptic peptides, numerous new, chromatographically separated peaks were observed (Fig 1A, bottom panel). Some of these new peaks had intense UV signals, such as those annotated as H9α, L13α, L13β, and H14γ in Fig 1. These peaks, however, were not identified as mAb-A peptides in the original database searching until nonspecific-cleavages rules were applied.

**Fig 1.**
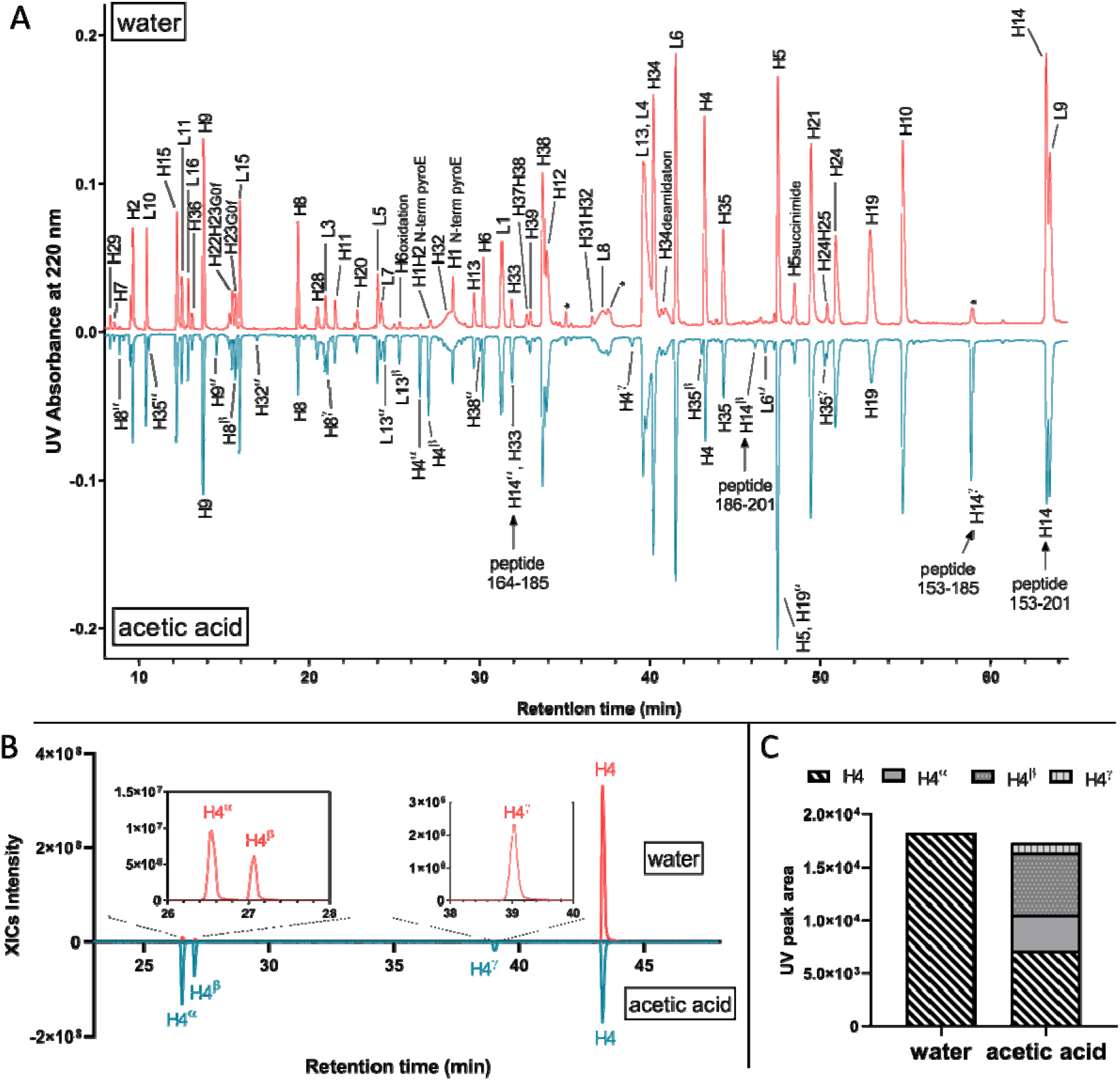
(A) Butterfly plot of UV chromatograms of mAb-A digested with Trypsin-1 reconstituted in HPLC-grade water and 50 mM acetic acid. Tryptic peptides are denoted using single letters (H = heavy chain; L = light chain) followed by the corresponding peptide number. Semitryptic and nontryptic peptides are denoted using the corresponding fully tryptic peptides that encompass their sequences, followed by Greek letters. Peaks annotated with asterisks are from autolysis. (B) Extracted ion chromatograms (XICs) of peptide H4 and the semitryptic peptides H4α, H4β, and H4γ. The abundances of semitryptic species were significantly higher when acetic acid was used for reconstitution. (C) UV peak integrals representing peptide H4 and the corresponding semitryptic peptides H4α, H4β, and H4γ showed that the sum of integrals between the two conditions were similar; however, H4α, H4β, and H4γ emerged in the acetic acid condition, at the cost of H4 signals.

Characterizations based on the MS/MS spectra indicated that these newly emerged peaks corresponded to semitryptic and nontryptic peptides of mAb-A. For instance, in addition to the CH1 domain H14 peptide (amino acid 153–201 with no missed cleavages), which eluted at min, we also identified peptides H14α, H14β, and H14γ, all of which carried nontryptic cleavages. H14γ (peptide 153–185) and H14β (peptide 186–201) were the two semitryptic counterparts that formed H14 via cleavage at Y185, whereas H14α (peptide 164–185) was a nontryptic peptide generated by simultaneous cleavages at W163 and Y185 (Fig 1A). The intensity of H14γ in condition 1 was nearly comparable to that of H14, indicating a strong preference to cleave at Y185 for this particular peptide when Trypsin-1 was reconstituted in 50 mM acetic acid instead of water. There were, however, many other peptides with adventitious, nonspecific cleavages. For example, the extracted ion chromatograms (XICs) of H4 and the corresponding peptides from nontryptic cleavages (H4α, H4β, H4γ) in both condition 1 and condition 2 (Fig 1B) demonstrated dramatic differences in signal abundances, such that peptides H4α, H4β, and H4γ, the signals of which were negligible in condition 2, grew significantly in condition 1. Interestingly, the signal of H4 decreased in condition 1 compared with condition 2, owing to the high yield of nontryptic cleavages. The UV peak areas of these peptides indicated an abundance decrease of ~60% for H4, with H4α, H4β, and H4γ emerging as new species in condition 1 (Fig 1C).

The incidence and extent of adventitious, nonspecific cleavages increased when Trypsin-1 was reconstituted in 50 mM acetic acid, producing more peaks in the chromatographic profile. Therefore, we further annotated the UV chromatogram to denote the visible new peaks (characterized as semitryptic and/or nontryptic peptides) by their corresponding fully tryptic peptides that encompass these sequences, using Greek letters in alphabetic order (α, β, γ, etc.; Fig 1A, bottom panel). Although it is noteworthy that all trypsins generated a certain amount of semitryptic and/or nontryptic cleavages [22, 26, 27], we observed that these adventitious, nonspecific cleavages increased markedly only when Trypsin-1 was reconstituted in a mildly acidic environment prior to digestion. The same phenomenon was repeated by using Trypsin-1 reconstituted in 1 mM HCl instead of 50 mM acetic acid (Supplementary Fig S1).

Our results indicate that acidic reconstitution of Trypsin-1 can significantly affect subsequent tryptic digestion by generating a greater extent of adventitious, nonspecific cleavages. In addition to antibodies, other molecule modalities, ranging from small proteins to adeno-associated virus (AAV) capsid proteins, were also accompanied by significant appearances of new peaks, owing to increased nontryptic activities induced by acidic reconstitution. Fig S2A shows tryptic digestion profiles (as TICs) of a 17-kDa protein with Trypsin-1 resuspended in HPLC-grade water and 1 mM HCl. The two visible new peaks in the 1 mM HCl condition were identified as peptide 30–48 and peptide 91–110, which resulted from W48 and Tyr110 cleavages, respectively.

### Time-dependent nontryptic activities

We found that the extent of the nontryptic activities induced by acidic reconstitution of Trypsin-1 increased with the length of the reconstitution period. Interestingly, when water was used for reconstitution, the level of nontryptic activities remained low.

As an example, samples of NISTmAb (an IgG1 antibody) were subjected to a 3.5-h tryptic digestion with Trypsin-1 reconstituted in 50 mM acetic acid at room temperature for five different time periods: t0, 1 h, 2 h, 4 h, and 6 h, where t0 corresponds to the immediate use of trypsin for digestion after dissolving. The control was Trypsin-1 reconstituted in HPLC-grade water at room temperature for the same periods.

To evaluate nontryptic activities as a function of trypsin reconstitution time, we monitored the signals of four NISTmAb semitryptic peptides, namely, heavy chain 151–183 and 184–213 as obtained from the cleavage at heavy chain Tyr183, light chain 61-86, and 87-102, as obtained from the cleavage at light chain Tyr 86, and their corresponding fully tryptic peptides, heavy chain 151–213 and light chain 61–102. We show in Fig 2A and 2B, using semitryptic peptide heavy chain 151-183 as an example, that the signal of this peptide markedly increased from t0 to 4 h with reconstitution in 50 mM acetic acid. The XIC of each peptide at each time point was extracted using monoisotopic mass, and the XIC integrals were plotted as a function of reconstitution time under the two reconstitution conditions (Fig 2C). For reconstitution with 50 mM acetic acid, an uptrend of XIC areas were seen for all four semitryptic peptides, whereas a downtrend was observed for the corresponding fully tryptic peptides. These observations strongly suggest that the increased abundances of semitryptic peptides occurred at the cost of fully tryptic peptide signals with a longer reconstitution time in acid. In contrast, with water reconstitution, the XIC areas of both semitryptic and fully tryptic peptides remained unchanged. It is noteworthy that the level of nontryptic activities remained low throughout the 6-h reconstitution period, as evidenced by the low abundance of all semitryptic peptides when water was used for reconstitution (Fig 2C).

**Fig 2.**
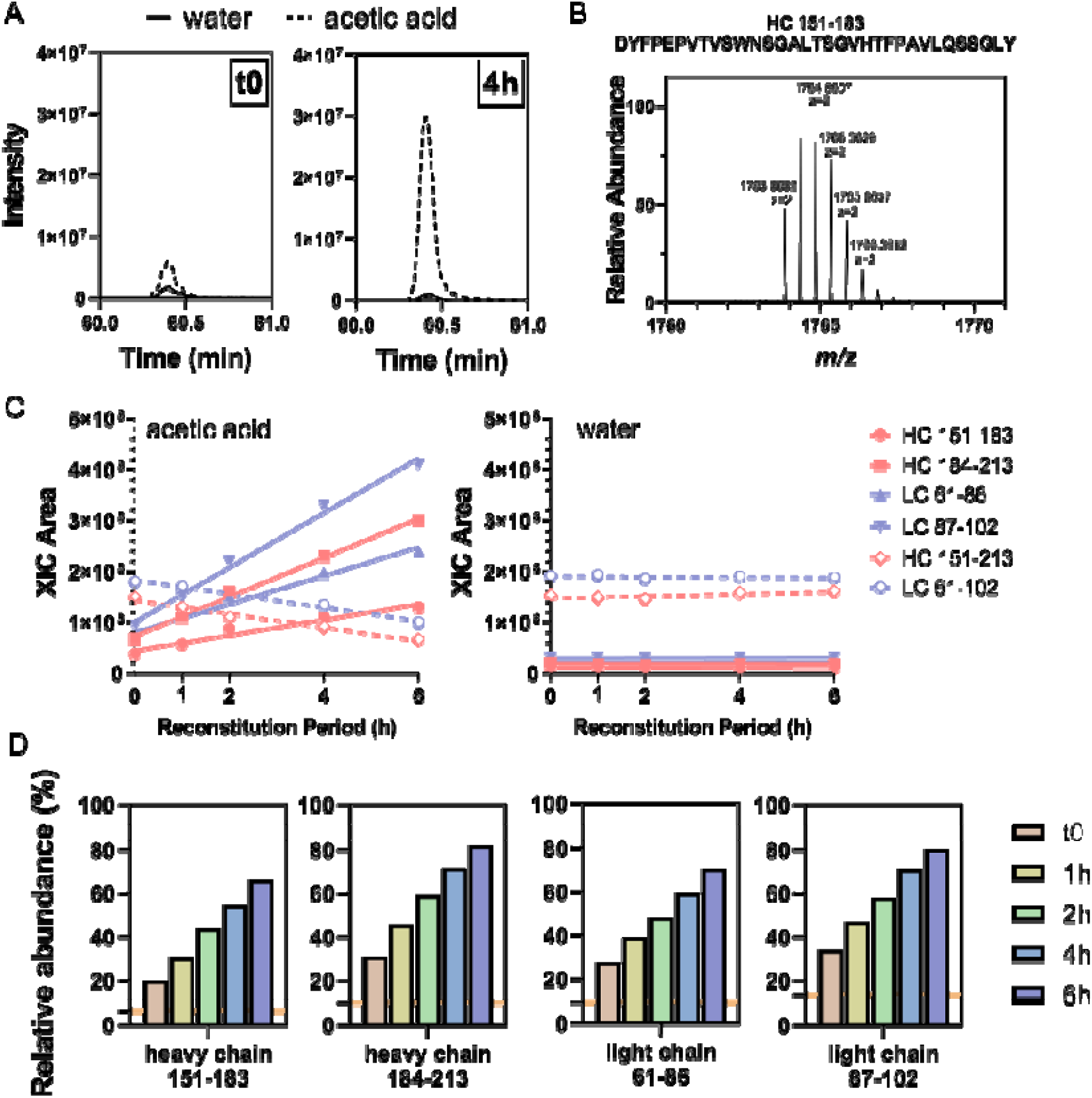
(A) XICs of NISTmAb peptide heavy chain 151–183 by monoisotopic mass (m/z 1763.8582) showed increased intensity with reconstitution in 50 mM acetic acid compared with reconstitution in water. The difference in XIC intensity between the two reconstitution conditions was more pronounced with the 4-h reconstitution period. Precursor-ion spectrum of NISTmAb peptide heavy chain 151–183 (z = 2), a semitryptic peptide generated from nontryptic cleavage at Y183. (C) When 50 mM acetic acid was used for Trypsin-1 reconstitution, all four semitryptic peptides showed increasing abundances with longer reconstitution times of up to 6 h, and the corresponding two fully tryptic peptides showed decreasing abundances. With water as the reconstitution condition, the abundances of all peptides remained unchanged, and those of the four semitryptic peptides were consistently low. (D) Abundances of the four semitryptic peptides relative to their corresponding fully tryptic peptides as a function of length of reconstitution period. Orange horizontal line indicates the corresponding averaged relative abundances of each peptide with reconstitution in water.

The abundances of the four monitored NISTmAb semitryptic peptides relative to their corresponding fully tryptic peptides are shown in Fig 2D. Time-dependent increases were observed with acidic reconstitution, and markedly lower relative abundances were found with water reconstitution. Taking heavy chain peptide 184-213 as example, the time-dependence of its relative abundance reported 30% at t0 and grew to 80% with 6-h reconstitution in acetic acid; whereas the relative abundance was stable around 10% throughout the reconstitution period in water. These results demonstrate that the abundances of these peptides are sensitive to different reconstitution conditions and that their relative abundances might serve as indicators of the extent of nonspecific cleavages. A pre-run of NISTmAb tryptic digestion with monitoring of relative abundances of these diagnostic peptides prior to running experiment samples should provide a quick evaluation of nontryptic activities.

These results suggest that, although the recommended reconstitution solution for Trypsin-1 is 50 mM acetic acid, nonspecific cleavages occurred under this condition and progressed as a function of the length of the reconstitution period. In contrast, using water for reconstitution resulted in reproducible tryptic performance with minimal nontryptic activities.

### Implementation of NPD

NPD, an indispensable component of the multi-attribute method (MAM) that debuted in 2015, is an emerging approach for nontargeted purity testing via binary comparison between a reference sample and an unknown [16, 17]. The use of advanced software algorithms to automatically align the chromatograms and identify any new peaks in the samples according to predefined peak selection criteria can have significant advantages over visual inspection of the profiles, especially when new peaks co-elute with an existing peak or the visible baseline starts to interfere with the profile of new peaks. We used NPD to capture the “changed” peaks, which were then subjected to extensive database searching with nonspecific cleavage rules and any number of missed cleavages. In doing so, we sought to leverage the identification of peptides induced by nontryptic cleavages due to acidic resuspension in order to generalize the altered tryptic cleavage pattern and understand the preferred sites for such nontryptic activities. Using NPD, we compared protein samples digested by Trypsin-1 reconstituted in acetic acid with the samples digested by Trypsin-1 reconstituted in HPLC-grade water.

As an example, in a binary comparison of NISTmAb tryptic digestion with Trypsin-1 reconstituted in 50 mM acetic acid for 6 h (test sample) and Trypsin-1 reconstituted in water (reference), a total of 121 species were designated as “new peaks” (Fig S3A). Some of these species were deconvoluted to identical masses and retention times, indicating that they had different charge states (and therefore different m/z values) attributed to the same peptide. A total of 58 masses were deconvoluted from the 121 species, and roughly 50% of these masses were in the range of 1,200–1,800 Da (Fig S3B). Characterizations of the 58 peptide species identified 52 semitryptic and 6 nontryptic NISTmAb peptides (Table 1). For each peptide, the fold-change value in abundance from the reference to the test sample (plotted in Fig S4) was calculated as:

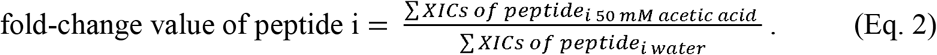

**Table 1.**
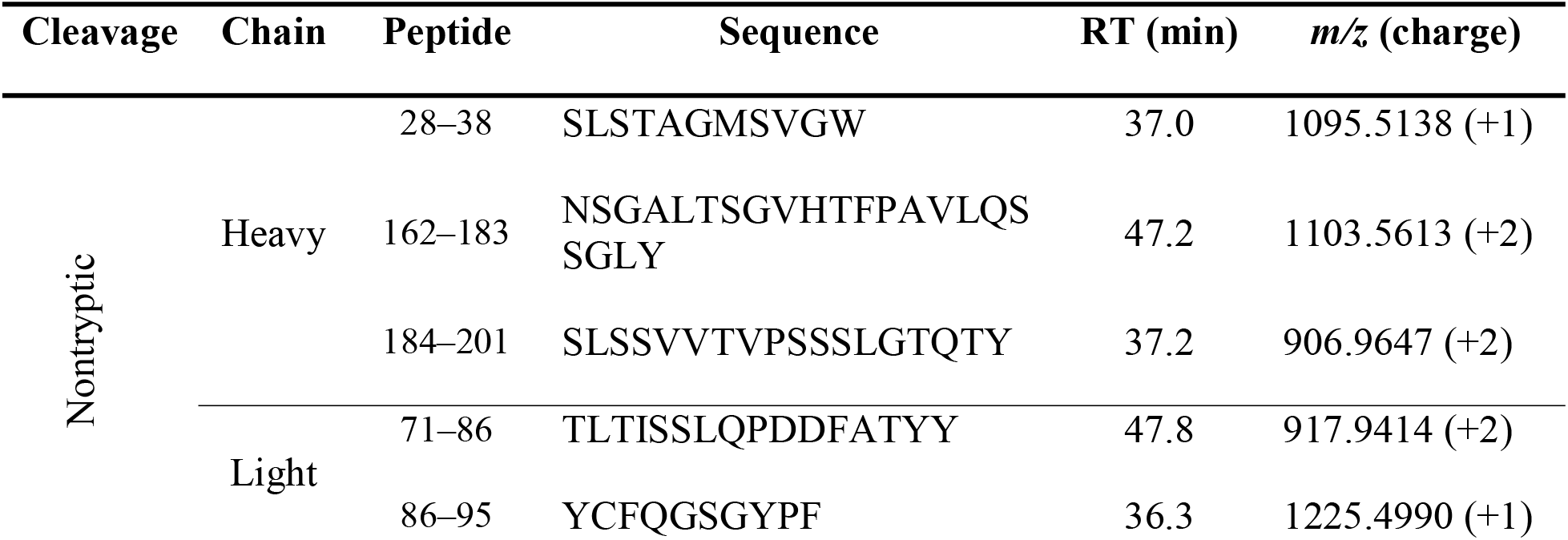
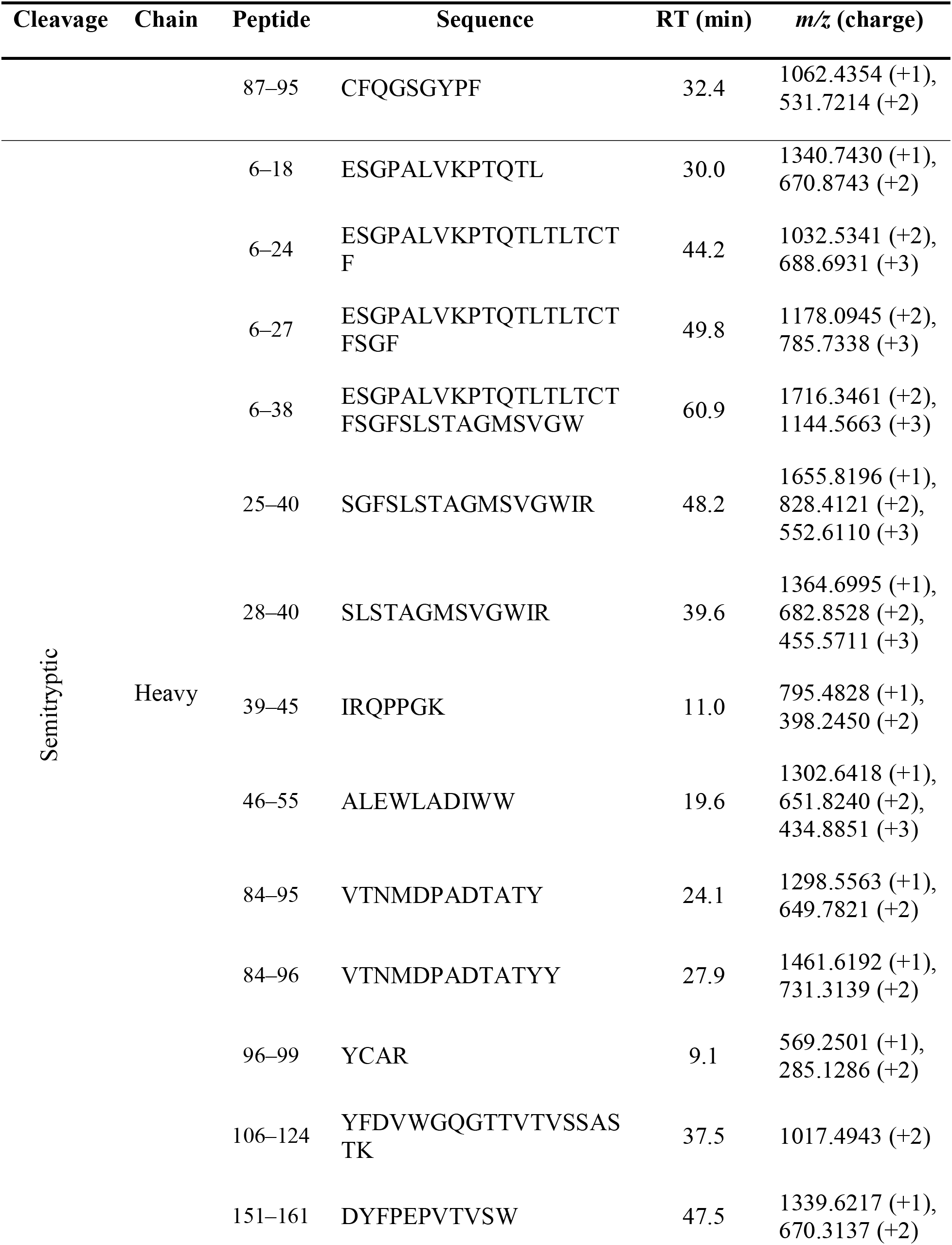
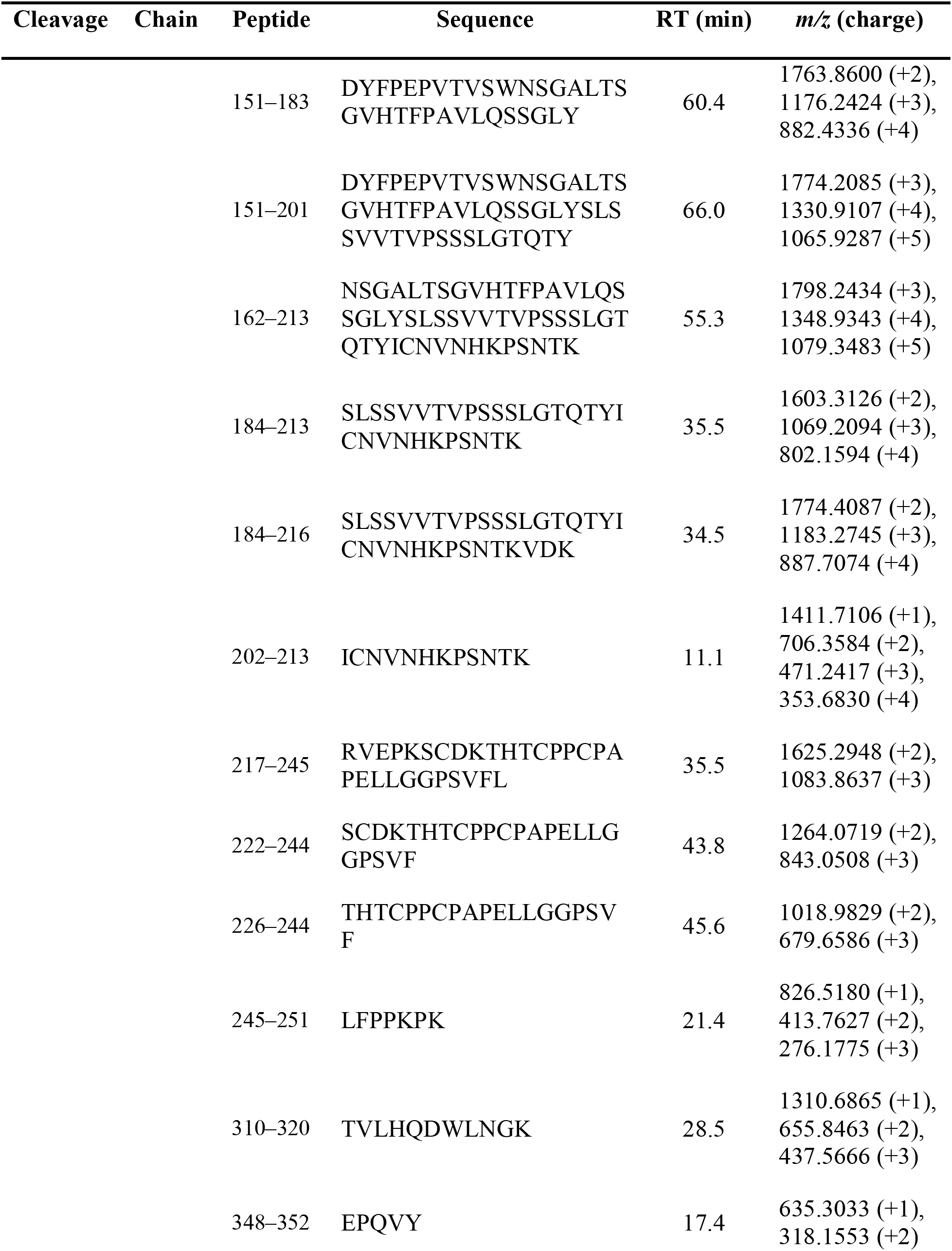
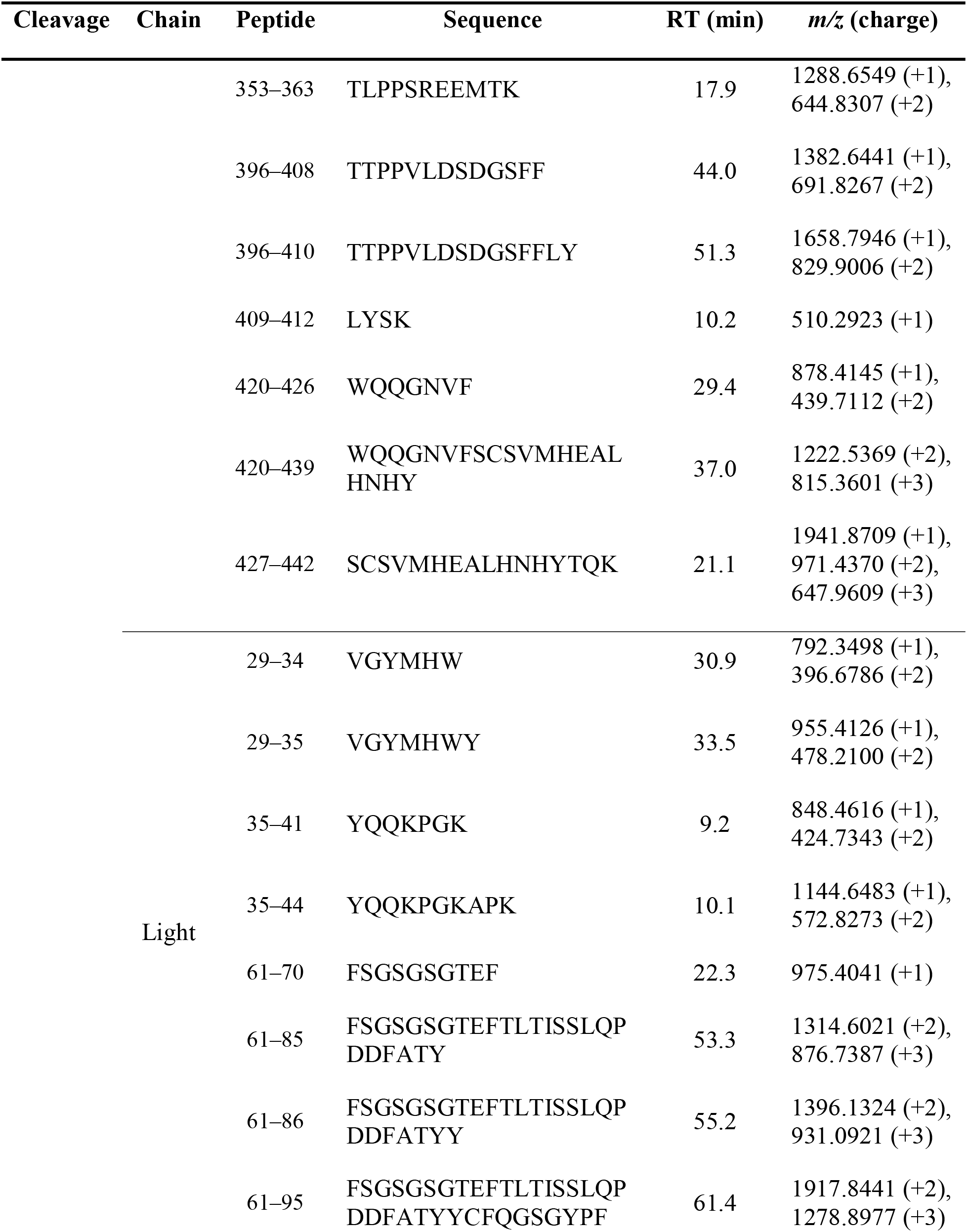
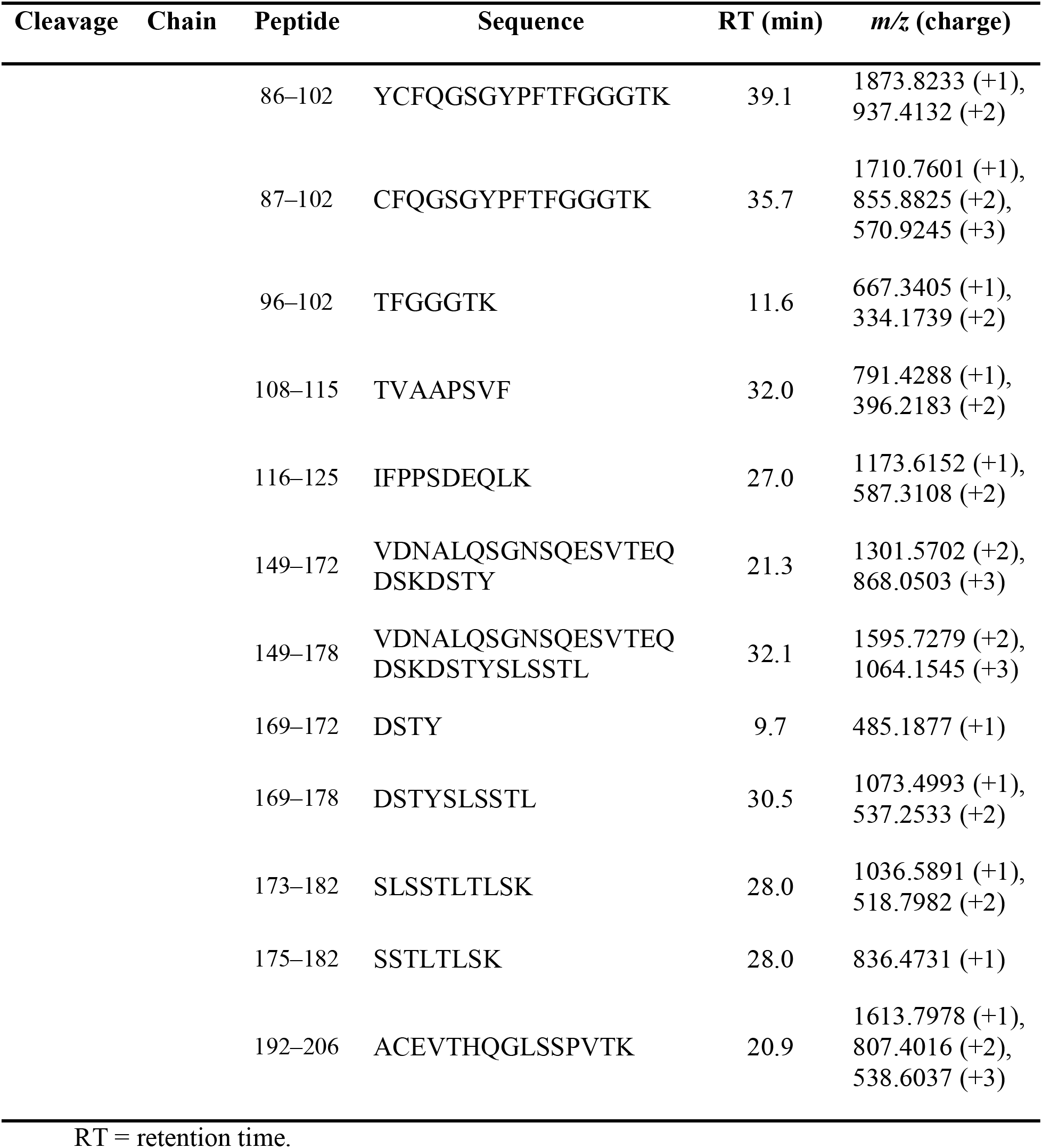
Identification of 6 nontryptic and 52 semitryptic NISTmAb peptides with their corresponding sequences, elution times, and *m/z* (charge)

All peptides demonstrated significant fold-change values (>3), consistent with the increased extent of nonspecific cleavages with the use of Trypsin-1 with acidic reconstitution. The fold-change values for semitryptic peptides ranged from 3 to 25, with a median of approximately 15, whereas those for nontryptic peptides were markedly higher, ranging from 100 to 300. These significant fold-change values indicate a considerable shift of Trypsin-1 cleavage specificity from highly tryptic (cleavages at R, K) to inclusion of some nontryptic sites.

With the registration of masses and elution times of NISTmAb nontryptic and semitryptic peptides by NPD analysis, we were able to provide a coarse-grained evaluation of the overall extent of nonspecific cleavages of Trypsin-1 as a function of reconstitution time. The XICs of the 58 NISTmAb peptides (52 semitryptic, 6 nontryptic) in each condition were summed together and divided by the XIC integrals of all identified peptides (per Eq. 1), giving the total fraction of peptides generated by nontryptic cleavages. With 50 mM acetic acid for Trypsin-1 reconstitution, the overall extent of nonspecific cleavages started at ~3.5% and further increased, at a rate of approximately 2.9% per hour, to as high as 22% when reconstitution reached 6 h in 50 mM acetic acid. The use of 1 mM HCl for reconstitution rendered a comparable extent of nonspecific cleavages (data not shown). In contrast, reconstitution in HPLC-grade water effectively inhibited the increase of nonspecific cleavages, the overall extent of which was consistently approximately 1% (Fig 3A). In addition to nonspecific cleavages, we also assessed the extent of missed cleavages, which ranged from 4% to 7% for all conditions (Fig S5A).

**Fig 3.**
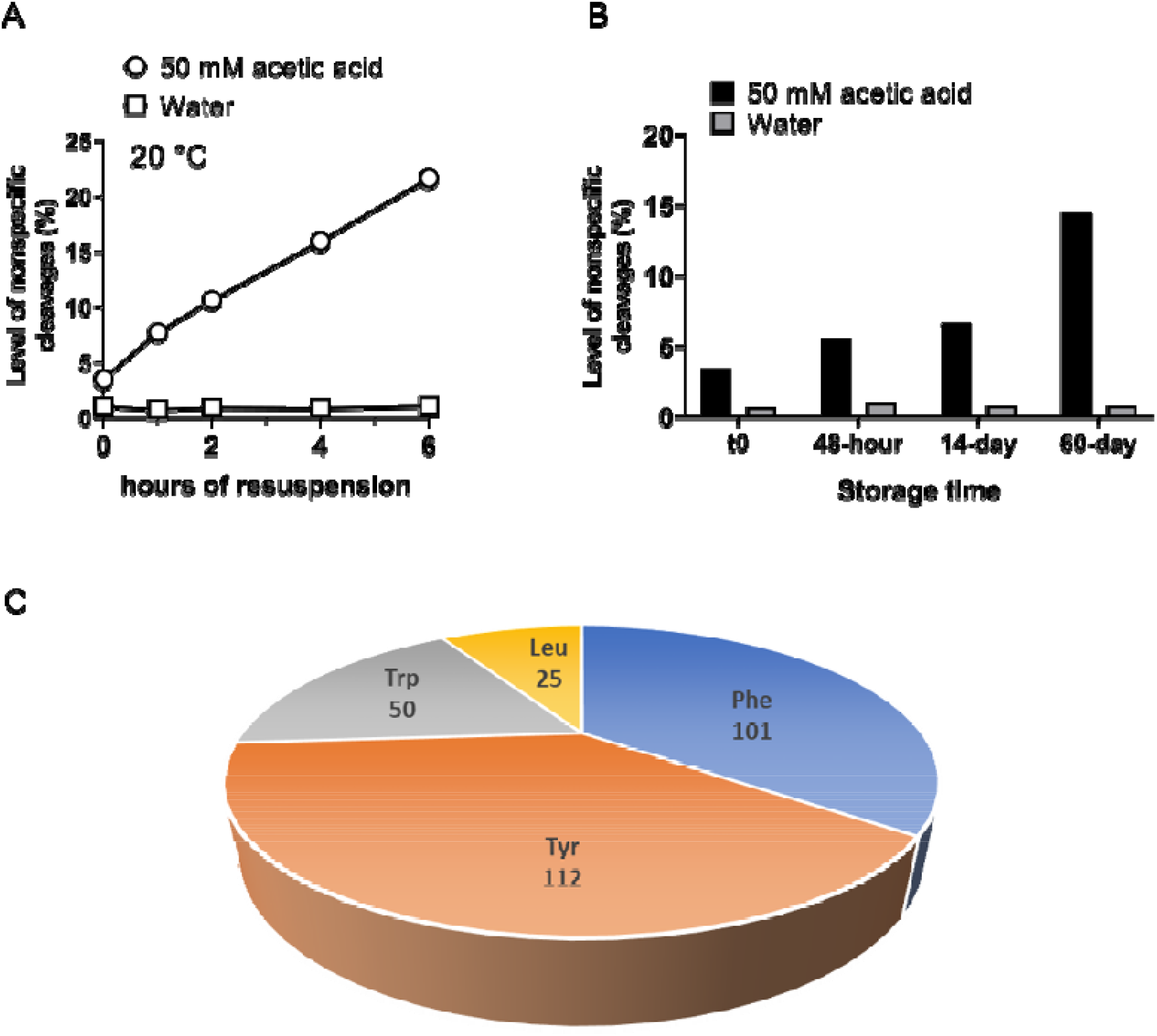
(A) Level of nonspecific cleavages as represented by the sum of XICs of the 52 NISTmAb semitryptic peptides and 6 nontryptic peptides relative to the sum of total identified peptides XICs as a function of trypsin resuspension time at room temperature (20°C). Dramatic differences were observed between the two conditions (i.e., 50 mM acetic acid and water); reconstitution in the acidic condition caused more nontryptic cleavages, the extent of which increased with longer reconstitution times. In contrast, the level of nonspecific cleavages remained consistent at approximately 1% with reconstitution of trypsin in water. (B) Evaluation of level of Trypsin-1 nonspecific cleavages under the recommended storage condition, with trypsin reconstituted in 50 mM acetic acid and stored at –80 °C for different periods, versus the storage condition using water for reconstitution. The yield of nonspecific cleavages under the recommended storage condition (50 mM acetic acid) was significantly higher and increased with the length of the storage period. (C) Demographic profile of amino acids involved in the semitryptic and nontryptic cleavages, based on 220 semitryptic peptides and 34 nontryptic peptides involving five biotherapeutic samples in addition to NISTmAb.

### Recommended storage conditions

The common recommendation for storing trypsins is in mildly acidic solution (e.g., 50 mM acetic acid) at low temperature. We investigated two sets of NISTmAb tryptic digestions, using Trypsin-1 reconstituted in HPLC-grade water and in 50 mM acetic acid. The trypsins were stored at −80°C for different periods before use (t0 and 2, 14, and 60 days). When the 60-day storage condition recommended by the vendor was used, the NISTmAb digests obtained from Trypsin-1 reconstituted in acetic acid had 15% total nonspecific cleavages. This was a noticeably higher level than that of digests obtained from Trypsin-1 reconstituted in water, which were consistently low (~1% of nonspecific cleavages) (Fig 3B). These results suggest a dramatically lower rate of increase in overall nontryptic activities when trypsin was subjected to low temperature with acidic reconstitution. However, the data also suggest that the low-temperature storage condition with Trypsin-1 reconstituted in water should be the optimal long-term storage condition with which to maintain the desired performance of trypsin, as evidenced by the minimal level of nonspecific cleavages. We did not observe an increase in the extent of missed cleavages during the long-term storage of Trypsin-1 (Figure S5B). Moreover, no impact on sequence coverage or PTM quantitation was observed for trypsin subjected to long-term storage and reconstitution in water (data not shown).

### Demographic profile

To populate the pool of semitryptic and nontryptic peptides caused by acidic reconstitution and/or storage of trypsin, NPD analysis of other biologics was performed. An investigation of preferred nonspecific cleavage sites was based on 220 semitryptic peptides and 34 nontryptic peptides, using five additional biotherapeutic samples besides NISTmAb. The demographic display of nonspecific cleavage sites based on these peptides indicated that four amino acids were accountable, namely, Tyr, Phe, Trp, and Leu. Approximately 90% of these cleavages occurred at the C-terminal of aromatic residues Tyr, Phe, and Trp, whereas Leu accounted for the remaining 10% (Figure 3B). This observation suggests that acidic reconstitution of Trypsin-1 leads to a shift in specificity, from highly specific for Lys and Arg to other amino acids, including Tyr, Phe, Trp, and Leu.

### Different vendors, different quality

In addition to Trypsin-1, we tested seven other commercial trypsins (Table 2) and assessed the extent of nontryptic activities and the effects of different reconstitution conditions.

**Table 2.** Overview of the eight commercial trypsins

| Name | Vendor | Catalog no. | Source | Recommended Reconstitution* |
| --- | --- | --- | --- | --- |
| Trypsin-1 | Promega | V5280 | Porcine | 50 mM acetic acid |
| Trypsin-2 | Promega | V5111 | Porcine | 50 mM acetic acid |
| Trypsin-3 | G-Biosciences | 786-245 | Porcine | 50 mM acetic acid |
| Trypsin-4 | G-Biosciences | 786-245B | Bovine | 50 mM acetic acid |
| Trypsin-5 | Princeton Separations | EN-151 | Porcine | Water |
| Trypsin-6 | Roche | 11418025001 | Bovine | 1% acetic acid or 1 mM HCl |
| Trypsin-7 | Pierce | 90057 | Porcine | 50 mM acetic acid |
| Trypsin-8 | Sigma | T-6567 | Porcine | 1 mM HCl |

\*Recommended reconstitution condition is based on product instruction of each trypsin

Lyophilized trypsin from each vendor was reconstituted in HPLC-grade water and in 50 mM acetic acid and kept at room temperature for 4 h before digestion of NISTmAb. The four diagnostic NISTmAb semitryptic peptides (heavy chain 151–183 and 184–213 and light chain 61–86 and 87–102) and the corresponding fully tryptic peptides (heavy chain 151–213 and light chain 61–102) were employed. The relative abundance of each peptide was calculated as the ratio of its XIC integral over its summed XIC integrals and the corresponding fully tryptic peptides. The results of all eight trypsins are summarized in Figure 4.

**Fig 4.**
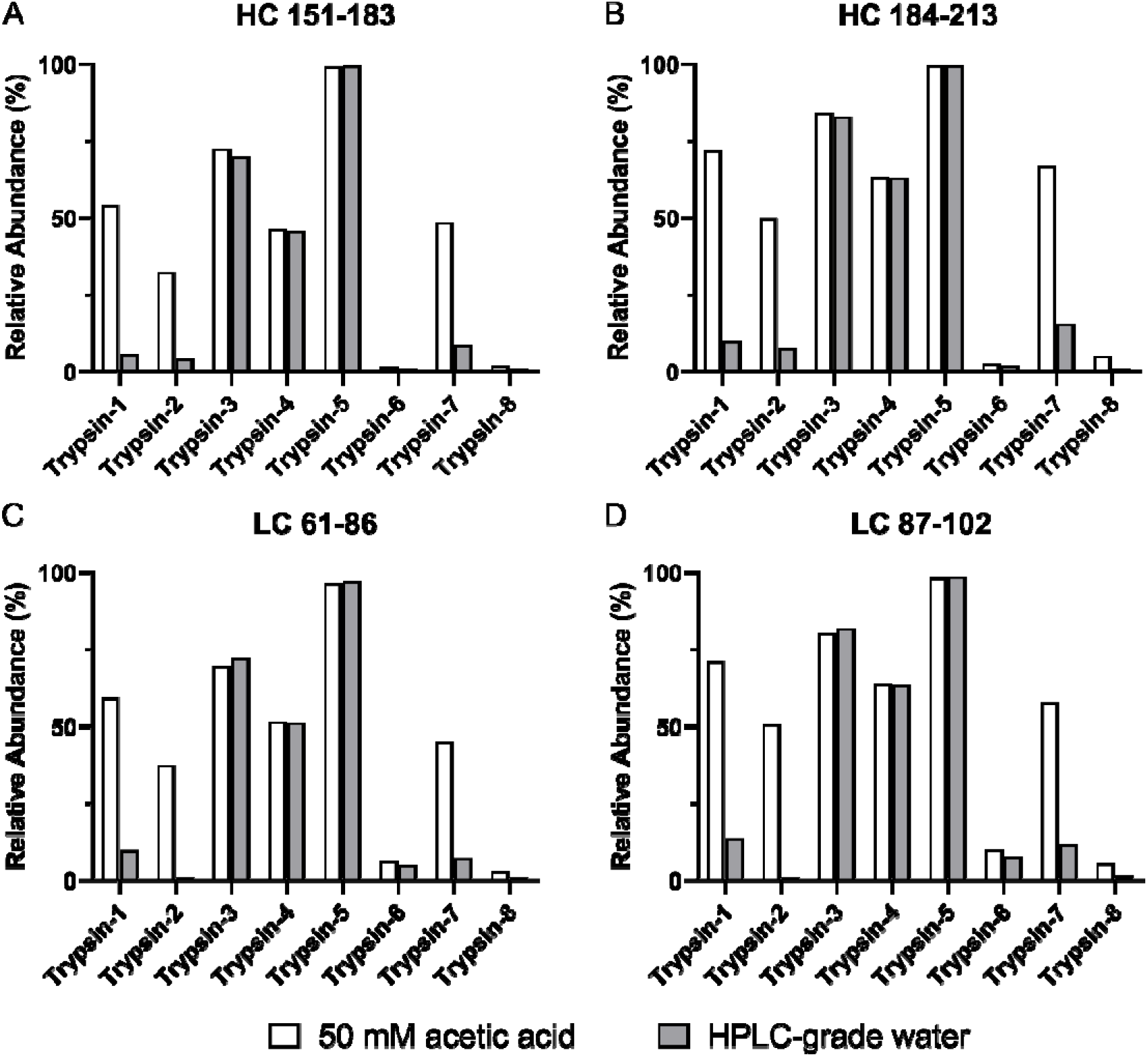
Relative abundance of selected NISTmAb semitryptic peptides for evaluation of nonspecific cleavages. Shown are (A) heavy chain 151–183, (B) heavy chain 184–213, light chain 61-86, and (D) light chain 87–102, as generated from tryptic digestion with eight commercial trypsins (Trypsin-1 to Trypsin-8). Each trypsin was reconstituted in 50 mM acetic acid (white bar) and HPLC-grade water (gray bar) at room temperature for 4 h before immediate use for digestion.

According to the manufacturers’ product information, all eight trypsins had been pretreated with TPCK and chemically modified and were claimed to afford high specificity.

Nevertheless, the level of nonspecific cleavages and the responses to the two reconstitution conditions by each trypsin were noticeably different. Trypsin-2 and Trypsin-7, much like Trypsin-1, showed an increased extent of nonspecific cleavages under the acetic acid reconstitution condition, but the level could be minimized by using water for reconstitution; although for these three trypsins, the vendors’ recommended reconstitution solvent is 50 mM acetic acid (Table 2). Nevertheless, not all trypsins were responsive to the different reconstitution conditions. Trypsin-3, Trypsin-4, and Trypsin-5 each showed comparable extents of nonspecific cleavages between the two reconstitution conditions, but the high percentage of semitryptic peptides indicated undesirable specificity regardless of the reconstitution condition. Two trypsins (Trypsin-6 and Trypsin-8) showed high specificity and consistently low levels of nonspecific cleavages under both reconstitution conditions. In addition, the results revealed the differing fidelity of trypsins from different manufacturers, which play a pivotal role in the distinct levels of specificity and responses to different reconstitution conditions.

### Possible causes of nontryptic activities

Our results suggest that different processes used for manufacturing of trypsins were accountable for the diverse nontryptic activities we observed, as trypsins from some vendors showed better specificity and tolerance to acidic reconstitution conditions than others. The observed nonspecific cleavages were unlikely to be due to chymotrypsin contamination, owing to the significant increase in the extent of nonspecific cleavages and their time dependence only when trypsin was subjected to acidic reconstitution. However, other contaminants from purification and chemical treatments might be possible. Another cause could be the formation of pseudotrypsin (Ψ-trypsin), a known variant of trypsin generated from the bond opening between K176 and D177 following an interchain split between K131 and S132 that yields α-trypsin [28, 29]. Coincident with our observations, pseudotrypsin also demonstrated a preference of cleavages after aromatic residues (Tyr, Phe, Trp) in addition to having characteristic trypsin properties [14, 30].

## Conclusions

Our results, with focuses on Trypsin-1, reveal a significantly increased level of nonspecific cleavages during the trypsin digestion process when trypsin is reconstituted or stored in a mildly acidic environment. In our investigation, the level of such nontryptic activities was proportional to the reconstitution/storage period. We demonstrated that the level of nonspecific cleavages, however, could be minimized to 1% simply by using HPLC-grade water for reconstitution. Besides Trypsin-1, several other commercial trypsins exhibited markedly compromised specificity when stored under conditions recommended by the manufacturer, potentially resulting in lack of reproducibility and sensitivity in LC-MS/MS– based research and applications. Based on our results, we recommend reevaluation of the recommended reconstitution of trypsins with 50 mM acetic acid. Our adoption of NPD analysis for the identification of semitryptic and nontryptic peptides enabled the demographic investigation of residues that were accountable for increased rates of nonspecific cleavages, whereby Tyr, Phe, Trp, and Leu were found to be the preferred sites involved in nontryptic activities.

## Supporting Information

**Fig S1.** Overlay of ultraviolet chromatograms of trypsin-digested monoclonal antibody A, using trypsin reconstituted in 1 mM HCl, 50 mM acetic acid, and high-performance liquid chromatography (HPLC)–grade water. The peak profiles of the digestion with trypsins reconstituted in acid were highly similar to, but different from, those with trypsin reconstituted in water. Nonspecific cleavages were significantly higher with acetic acid reconstitution. The dashed-line boxes indicate selected regions in which additional peaks corresponding to nonspecific cleavages arose.

**Fig S2.** (A) Total ion current chromatograms corresponding to the tryptic digestion of a 17-kDa protein with trypsin reconstituted in HPLC-grade water (upper panel) and 1 mM HCl (lower panel). The two visible new peaks (shaded in blue) were identified as peptides 30–48 and 91–110 from nontryptic cleavages at W48 and Tyr110, respectively. (B) Extracted ion chromatograms of peptides 30–48 and 91–110, showing the dramatic increase in nontryptic cleavages that occurred when 1 mM HCl was used for trypsin reconstitution.

**Fig S3.** (A) New peak detection analysis designated 121 species as “new,” based on the predefined peak selection criteria. The apex retention time of each species versus the corresponding monoisotopic m/z is plotted. (B) Mass distribution of 58 peptides deconvoluted from the 121 species. Approximately 50% of peptides had masses ranging from 1,200 to 1,800 Da.

**Fig S4.** Box plot showing the fold-change values in abundance for all identified semitryptic and nontryptic peptides. Although all peptides showed fold-change values of a minimum of 3, nontryptic peptides demonstrated more significant fold-change values than did semitryptic peptides.

**Fig S5.** Assessment of the extent of missed cleavages showed that the level of missed cleavages ranged from 4% to 7% for all conditions.

## Acknowledgments

The authors thank Chunlei Wang, Xiaoyu Chen, Sophie Inman, Jared Delmar, Meagan Prophet, Kevin Wons for providing input and discussion in this work. Editorial support was provided by Deborah Shuman of AstraZeneca.

## Author contributions

### Sample treatment and preparation

Ben Niu, Michael Martinelli II, Yang Jiao

### Data curation

Ben Niu

### Investigation and discussion

Ben Niu, Michael Martinelli II, Yang Jiao, Jihong Wang, Mingyan Cao, Eric Meinke

### Writing

Ben Niu

