## Supporting Information for "Nonspecific Cleavages Arising from Reconstitution of Trypsin under Mildly Acidic Conditions"

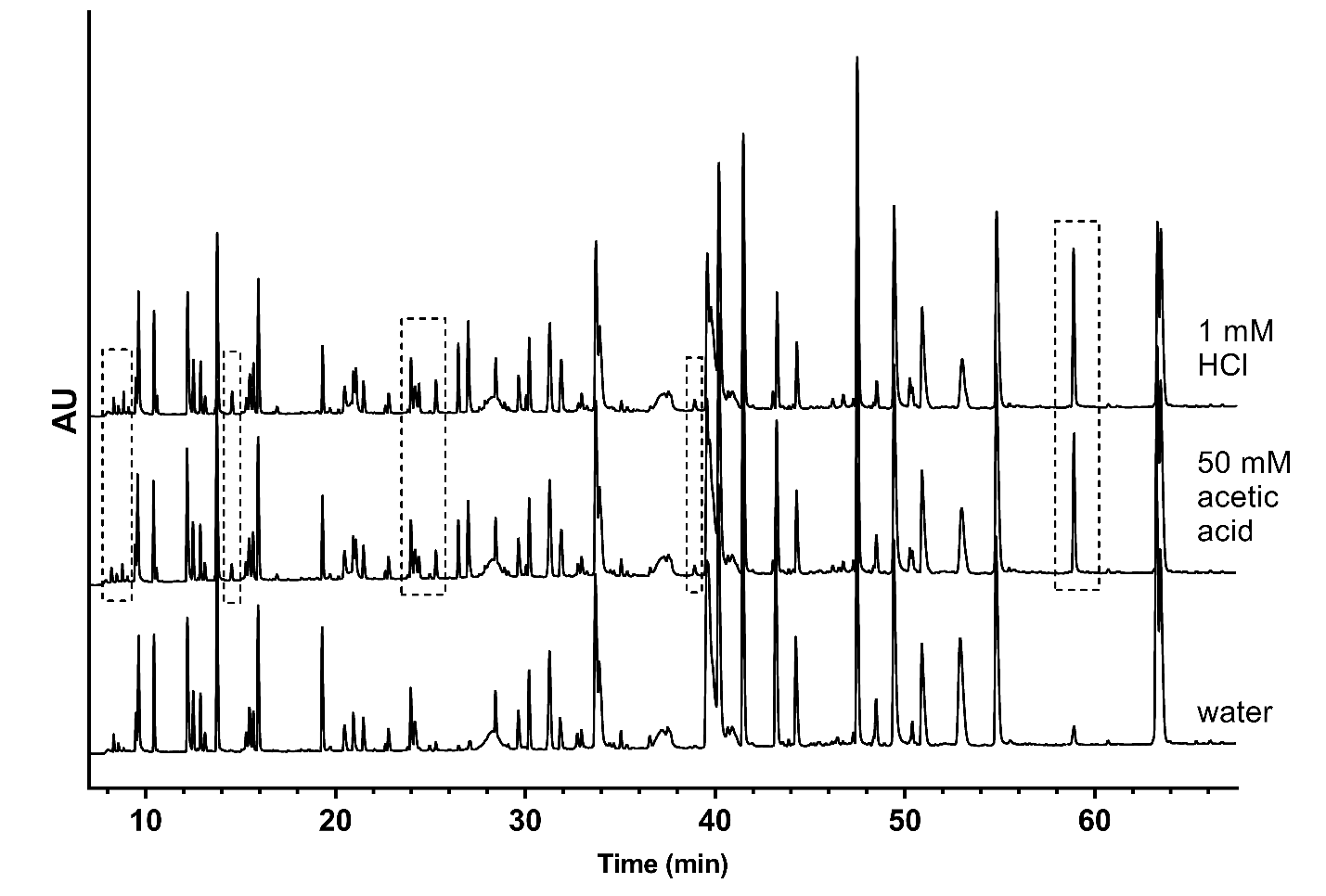

**Fig S1.** Overlay of ultraviolet chromatograms of trypsin-digested monoclonal antibody A, using trypsin reconstituted in 1 mM HCl, 50 mM acetic acid, and high-performance liquid chromatography (HPLC)–grade water. The peak profiles of the digestion with trypsins reconstituted in acid were highly similar to, but different from, those with trypsin reconstituted in water. Nonspecific cleavages were significantly higher with acetic acid reconstitution. The dashed-line boxes indicate selected regions in which additional peaks corresponding to nonspecific cleavages arose.

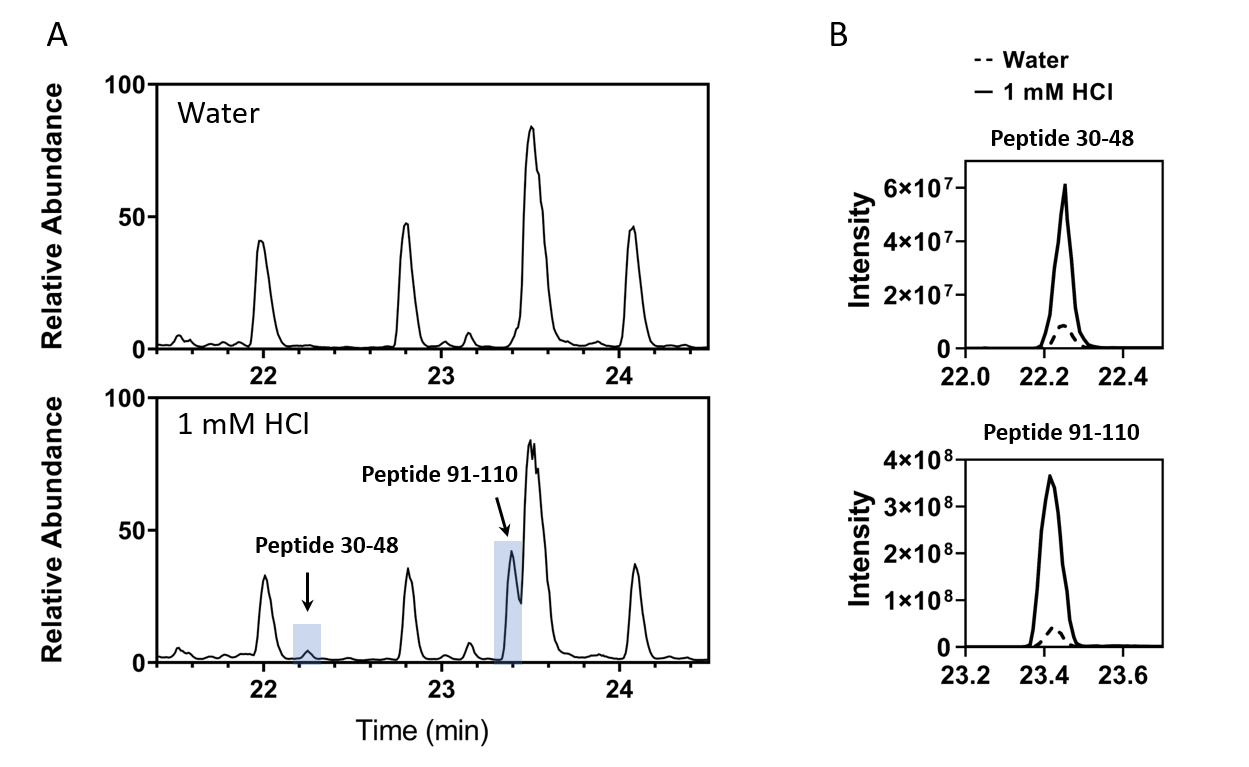

**Fig S2.** (A) Total ion current chromatograms corresponding to the tryptic digestion of a 17-kDa protein with trypsin reconstituted in HPLC-grade water (upper panel) and 1 mM HCl (lower panel). The two visible new peaks (shaded in blue) were identified as peptides 30–48 and 91–110 from nontryptic cleavages at W48 and Tyr110, respectively. (B) Extracted ion chromatograms of peptides 30–48 and 91–110, showing the dramatic increase in nontryptic cleavages that occurred when 1 mM HCl was used for trypsin reconstitution.

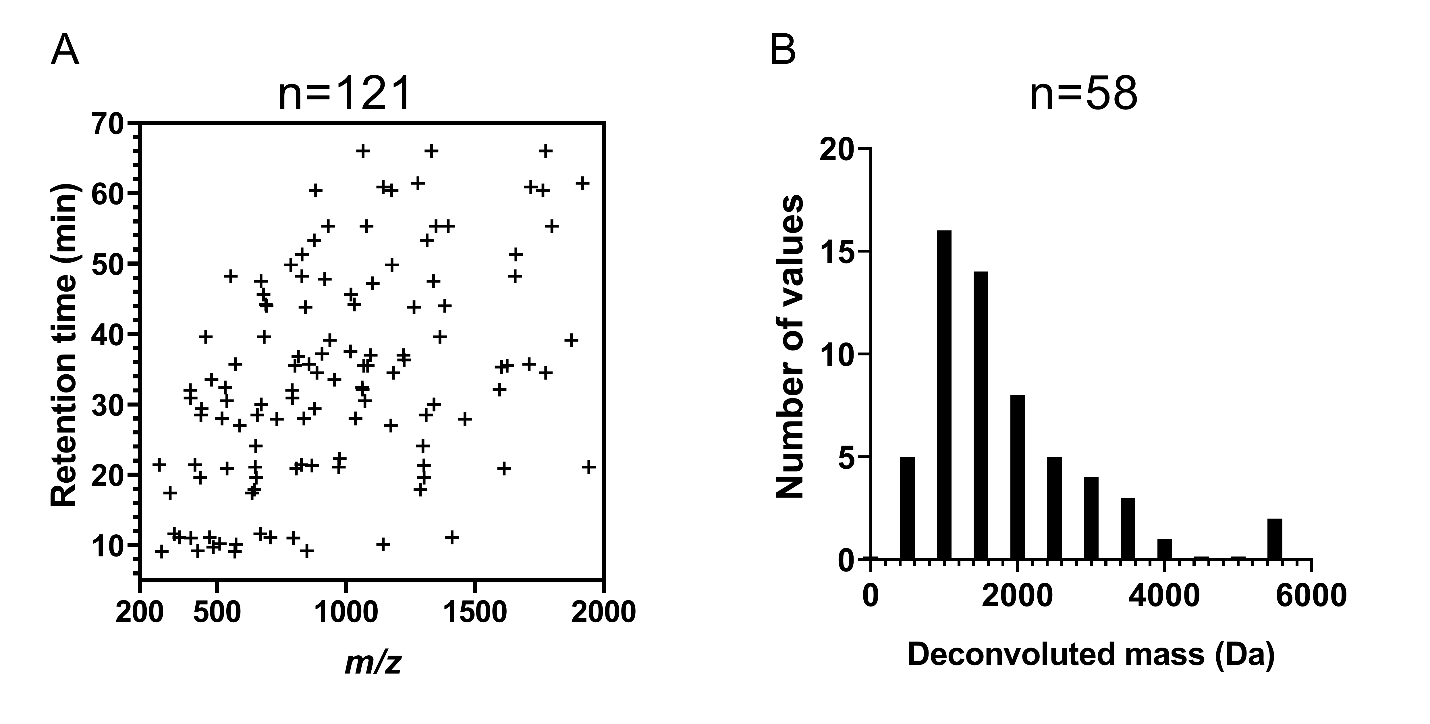

**Fig S3.** (A) New peak detection analysis designated 121 species as “new,” based on the predefined peak selection criteria. The apex retention time of each species versus the corresponding monoisotopic m/z is plotted. (B) Mass distribution of 58 peptides deconvoluted from the 121 species. Approximately 50% of peptides had masses ranging from 1,200 to 1,800 Da.

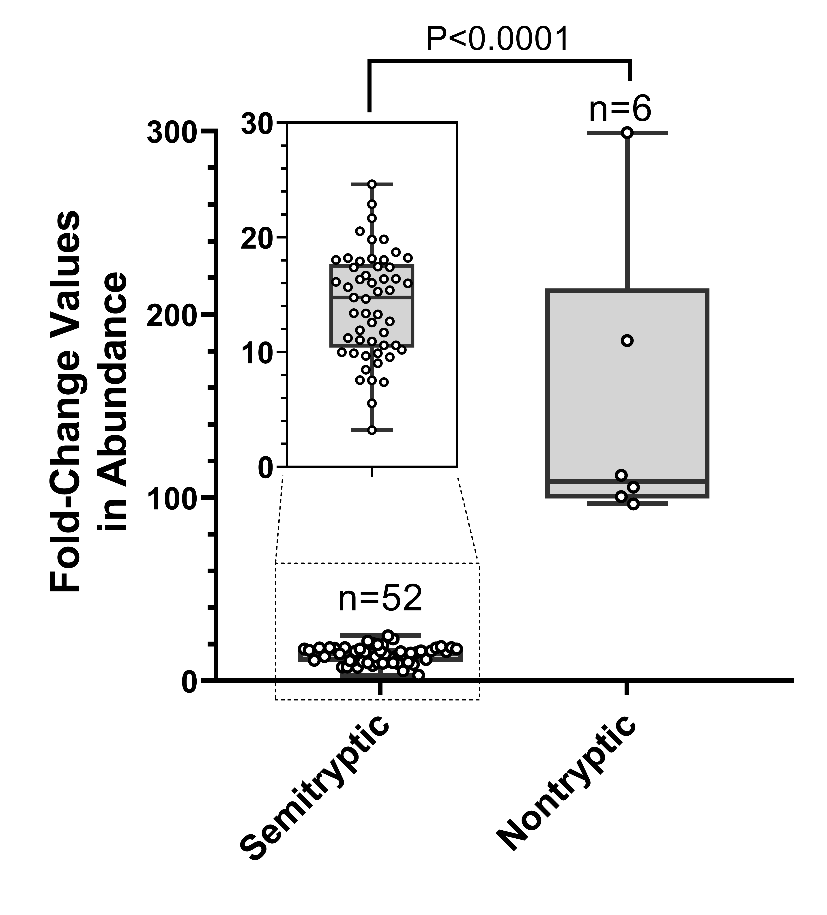

**Fig S4.** Box plot showing the fold-change values in abundance for all identified semitryptic and nontryptic peptides. Although all peptides showed fold-change values of a minimum of 3, nontryptic peptides demonstrated more significant fold-change values than did semitryptic peptides. Statistical analysis was performed using the Student *t* test.

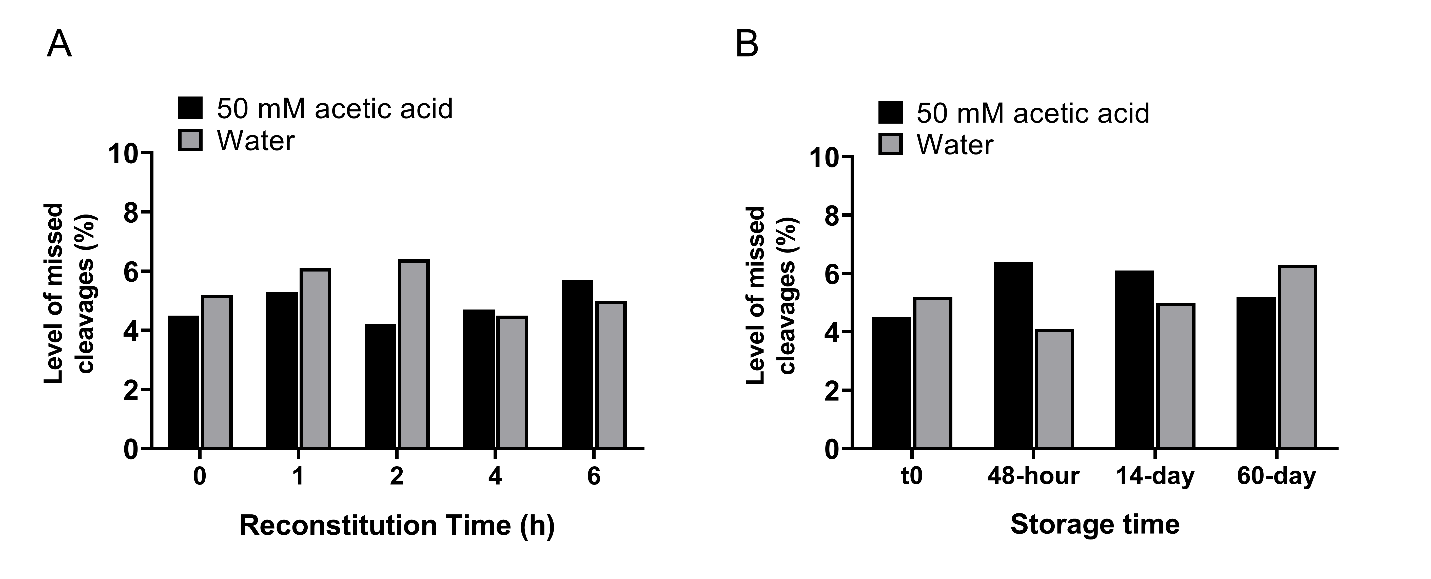

**Fig S5.** Assessment of the extent of missed cleavages showed that the level of missed cleavages ranged from 4% to 7% for all conditions.
